# A conserved machinery underlies the synthesis of a chitosan layer in the *Candida* chlamydospore cell wall

**DOI:** 10.1101/2021.01.19.427374

**Authors:** Leo D. Bemena, Kyunghun Min, James B. Konopka, Aaron M. Neiman

## Abstract

The polysaccharide chitosan is found in the cell wall of specific cell types in a variety of fungal species where it contributes to stress resistance, or in pathogenic fungi, virulence. Under certain growth conditions, the pathogenic yeast *Candida dubliniensis* forms a cell type termed a chlamydosospore, which has an additional internal layer in its cell wall as compared to hyphal or yeast cell types. We report that this internal layer of the chlamydospore wall is rich in chitosan. The ascospore wall of *Saccharomyces cerevisiae* also has a distinct chitosan layer. As in *S. cerevisiae*, formation of the chitosan layer in the *C. dublinien*sis wall requires the chitin synthase *CHS3* and the chitin deacetylase *CDA2*. In addition, three lipid droplet-localized proteins Rrt8, Srt1, and Mum3, identified in *S. cerevisiae* as important for chitosan layer assembly in the ascospore wall, are required for the formation of the chitosan layer of the chlamydospore wall in *C. dubliniensis*. These results reveal that a conserved machinery is required for the synthesis of a distinct chitosan layer in the walls of these two yeasts and may be generally important for incorporation of chitosan into fungal walls.

**Importance:** The cell wall is the interface between the fungal cell and its environment and disruption of cell wall assembly is an effective strategy for antifungal therapies. Therefore, a detailed understanding of how cell walls form is critical to identify potential drug targets and develop therapeutic strategies. This work shows that a set of genes required for assembly of a chitosan layer in the cell wall of *S. cerevisiae* is also necessary for chitosan formation in a different cell type in a different yeast, *C. dubliniensis*. Because chitosan incorporation into the cell wall can be important for virulence, the conservation of this pathway suggests possible new targets for antifungals aimed at disrupting cell wall function.

## Introduction

The cell wall is the interface between the fungal cell and the environment (1). In pathogenic fungi, the cell wall is critical for virulence as it mediates interactions with, and evasion of, the host immune system (2). Fungal cell walls are essential for viability and are a common target of antifungal drugs (3-6). Therefore, understanding the structure and assembly of the fungal wall is important for the development of antifungal therapies.

Fungal cell walls are composed primarily of heavily mannosylated proteins (referred to as mannan) and polysaccharides (1). In particular beta 1,3 glucans and chitin, a beta 1,4-*N*-acetylglucosamine polymer, are common structural components of fungal cell walls (1, 7, 8). Chitosan, a beta 1,4-glucosamine polymer created by deacetylation of chitin, is also found in fungal cell walls but is often limited to specific cell types or developmental stages (9-12). The presence of chitosan in cell walls can be critical for the organism. For example, in the pathogen *Cryptococcus neoformans*, chitosan in the wall dampens the host inflammatory response, and *Cryptococcus* strains unable to synthesize chitosan are avirulent (13-15). Chitosan is often found in conjunction with polyphenolic compounds, which has led to the proposal that chitosan-polyphenol complexes are a conserved architectural motif in fungal walls (16).

How chitosan is incorporated into the cell wall is not yet well understood. This process has been best-studied in the budding yeast, *Saccharomyces cerevisiae*, where chitosan is found uniquely in the walls of ascospores, a dormant cell type produced after meiosis by a process termed sporulation (17, 18). The ascospore wall consists of four distinct layers, named for their primary constituents, that are deposited in a sequential manner: mannan, glucan, chitosan and dityrosine (10, 19-22). The mannan and glucan layers form the inner layers of the ascospore wall and are similar in composition to layers in the vegetative cell wall (21). The outer ascospore wall, containing a layer of chitosan and a layer of the polyphenol dityrosine is unique to ascospores and confers resistance against environmental insults (10, 23, 24).

The chitin in the vegetative cell wall of *S. cerevisiae* is produced by three different chitin synthases, Chs1, 2 and 3 (25-27). However, during sporulation chitin is produced exclusively by Chs3 (28). Chitosan is generated when acetyl groups on chitin are removed by the sporulation-specific deacetylases, Cda1 and Cda2 (11, 29). Deletion of both *CDA1* and *CDA2* results in spore walls that contain chitin, but lack the chitosan layer. In addition, while the mannan and beta-glucan layers are present, the dityrosine layer is missing. Chitosan is therefore necessary for the formation of both layers of the outer cell wall (29). In contrast, formation of the chitosan layer is independent of the formation of dityrosine. Dityrosine is synthesized from L-tyrosine in the spore cytosol by the sequential action of the Dit1 and Dit2 enzymes (30) and mutants in either *DIT1* or *DIT2* result in loss of the dityrosine without any obvious effect on the chitosan layer (23).

In addition to the genes directly involved in chitosan or dityrosine synthesis, several other genes are required for the formation of one or more layers of the outer spore wall (31-35). Genes of unknown function such as *MUM3* and *OSW1*, as well as the cis-prenyltransferase encoded by *SRT1*, lack both the chitosan and dityrosine layers (34). In an *srt1*∆ mutant, Chs3 activity is reduced, suggesting that Srt1 contributes to spore wall formation through regulation of Chs3 (34). Srt1 is localized to a class of lipid droplets that is physically associated with the developing spore wall (34, 36). Mutants in the paralogous genes *LDS1, LDS2*, and *RRT8*, which encode lipid droplet-localized proteins, are specifically defective in the dityrosine layer (35). Whether the genes required for chitosan layer formation in *S. cerevisiae* are functionally conserved in other fungi has not been reported.

The human fungal pathogen, *Candida albicans* and its close relative, *Candida dubliniensis*, exhibit cell types with varying morphologies (37, 38). Though these *Candida* species are not known to produce ascospores, under certain conditions they produce a distinct, thick-walled cell type at hyphal tips termed a chlamydospore (37, 39). Chlamydospores are large round cells that are the result of mitotic divisions, unlike ascospores which package the haploid products of meiosis. The function of chlamydospores in the *Candida* life cycle is unknown. Nutrient limitation or low oxygen conditions are often required to induce the appearance of chlamydospores, and *C. dubliniensis* appears to undergo chlamydosporulation more readily than *C. albicans* (40, 41).

Ultrastructural studies revealed that the chlamydospore wall is more extensive than the wall of budding or hyphal *C. dubliensis* cells with an internal layer not found in those cell types (42). The structure and composition of this layer has not been well characterized. In the present study, we investigated the organization of the chlamydospore wall in *C. dubliniensis*. This work demnostrated that the unique internal layer of the chlamydospore wall is composed of chitosan. Moreover, genes encoding orthologs of *S. cerevisiae* proteins necessary for chitosan layer synthesis in ascospores are also required chlamydospore wall assembly. These results reveal that a conserved pathway underlies chitosan synthesis and incorporation in these two yeasts.

## Results

### *C. dubliniensis* forms chlamydospores on solid medium containing non-fermentable carbon sources

In examining the growth of clinical isolates of *C. dubliniensis* we discovered that growth on certain carbon sources induced chlamydospore formation. While chlamydospores were not observed in cultures grown on synthetic medium containing glucose or galactose, growth on N-acetyl glucosamine, glucosamine, glycerol, or acetate all led to hyphal growth and the appearance of chlamydospores (Figure 1). Three different clinical isolates of *C. dubliniensis* as well as the established *C. dubliniensis* strain SN90 (43) displayed this behavior, whereas *C. albicans* did not form chlamydospores on any of these media (K. M, unpublished obs.). Solid glycerol medium was particularly efficient at inducing chlamydospores (no chlamydospores were seen in liquid medium with any carbon source) (Figure 1). We took advantage of these induction conditions to examine the properties of the chlamydospore wall in *C. dubliniensis*.

**Figure 1.**
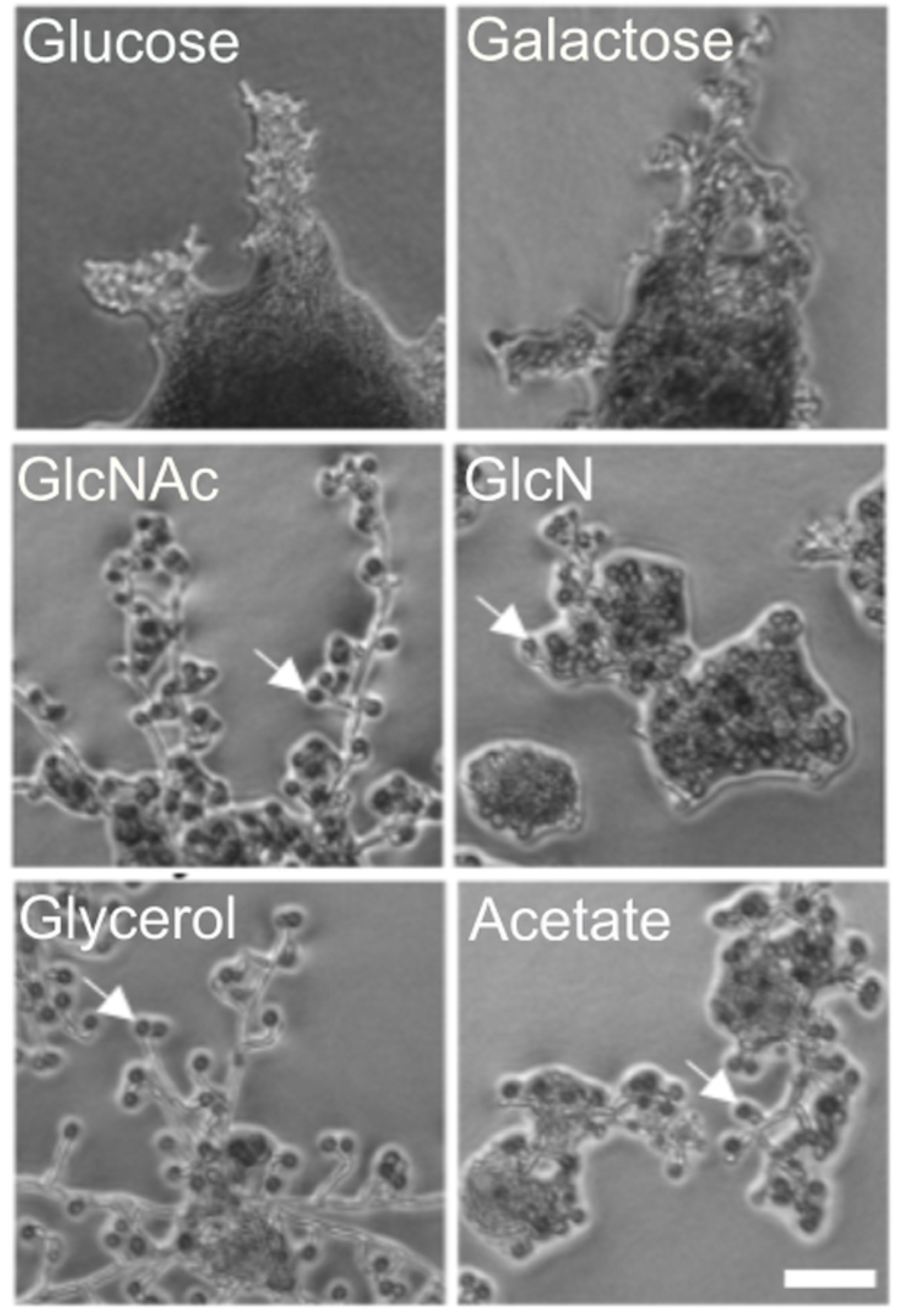
Effect of different carbon sources on the chlamydospore formation. A wild-type *C. dubliniensi*s strain (Cd1465) was spotted on synthetic agar medium containing the indicated carbon sources and were photographed on agar after 24 hours of growth. Gal-Galactose; GlcNAc – N-Acetylglucosamine; GlcN-Glucosamine. White arrows highlight examples of clamydospores. Scale Bar = 50 nm.

### The chlamydospore wall of *C. dubliniensis* contains chitosan but not dityrosine

In the *S. cerevisiae* ascospore wall, the dityrosine layer is underlain by a layer of chitosan and chitosan is found in association with polyphenol components in other fungal cell walls (9). The observation that chlamydospore walls of *C. albicans* contain dityrosine suggested that chlamydospore walls might contain chitosan as well (44). Chitosan can be specifically visualized using the stain Eosin Y, which has affinity for chitosan but not chitin (9, 35). When *C. dubliniensis* chlamydospores were stained with Eosin Y and examined by fluorescence microscopy, bright Eosin Y-dependent fluorescence was visible at the periphery of the chlamydospore (Figure 2A, B). The fluorescent signal was not observed on hyphal cells, consistent with the presence of chitosan specifically in the chlamydospore wall. Similar staining of *C. albicans* chlamydospores with Eosin Y has recently been reported (45).

**Figure 2.**
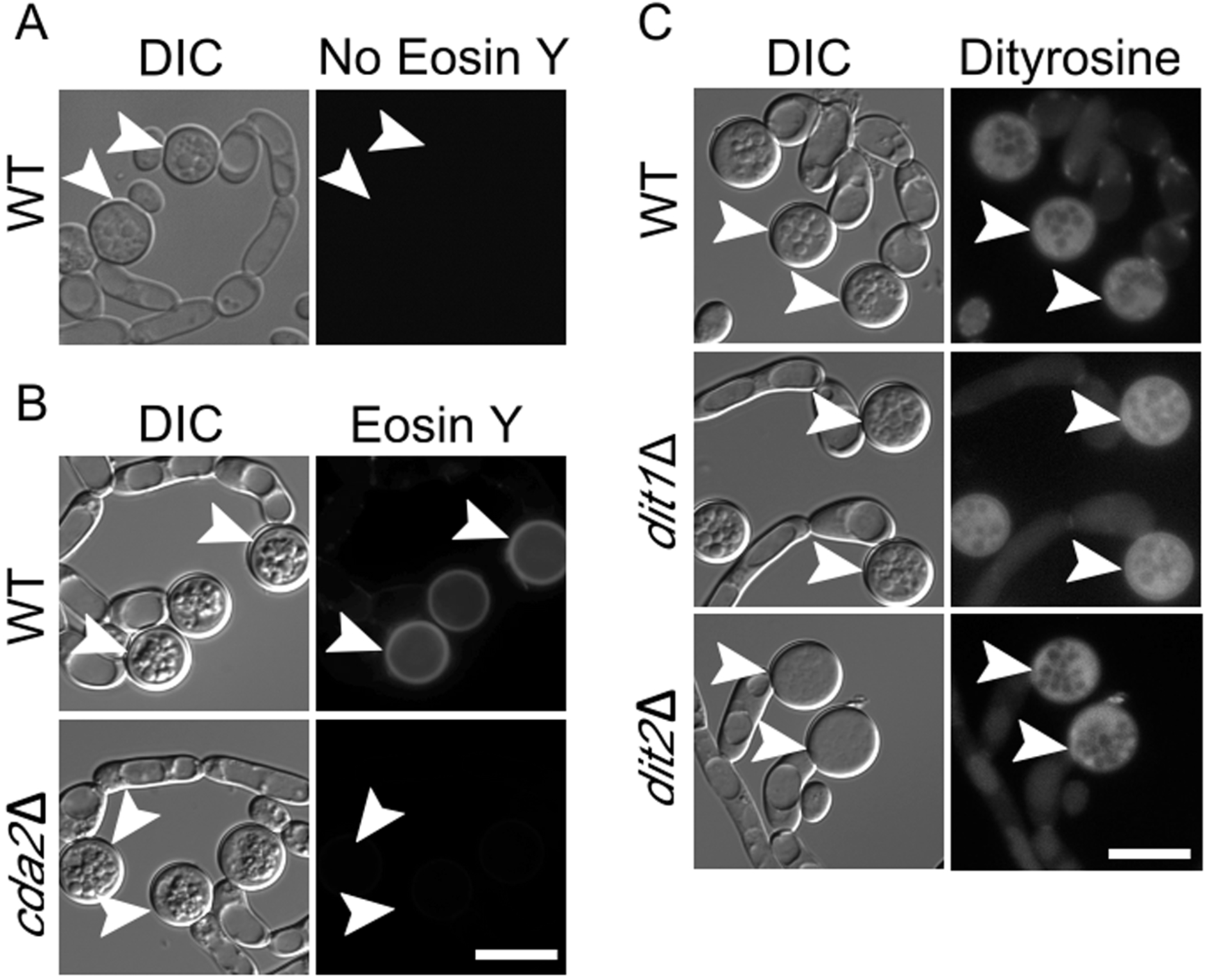
Fluorescence analysis of the chlamydospore wall of *C. dubliniensis*. (A) WT (Cd1465), *dit1*Δ (BEM9), or *dit2*∆ (BEM10) were grown on SG medium to induce chlamydospores and then visualized by differential interference contrast (DIC) or fluorescence microscopy using a dityrosine filter set (Ex. 320nm Em. 410nm). (B) Chlamydospores of WT (Cd1465) and *cda2*Δ (BEM7) strains were stained with Eosin Y to visualize the chitosan layer and imaged using a GFP filter set. WT chlamydospores with no Eosin Y staining are shown as control. Arrowheads indicate examples of chlamydospores visible in the images. Scale bar = 10µm.

To prove whether Eosin Y staining was specifically detecting chitosan, a genetic approach was used. The *C. dubliniensis* genome encodes one member of the chitin deacetylase enzyme family, Cda2 (Cd36_25340), required to convert chitin to chitosan. If Eosin Y staining is due to the presence of chitosan in the chlamydospore wall, this staining should be reduced or absent in a *cda2* deletion that lacks chitin deacetylase activity (9, 35).

*C. dubliniensis* is a diploid organism. To generate a *cda2*∆/*cda2*∆ deletion strain in *C. dubliniensis*, we utilized a transient CRISPR-Cas9 system originally developed for *C. albicans* (46). Double strand breaks in the two *CDA2* alleles were generated by transformation with two separate linear DNA fragments encoding a Cas9 enzyme and a guide RNA specific to a sequence within *CDA2*, respectively. In addition, the transformation included a “healing fragment” comprised of the *SAT1* cassette (47) flanked by 20 bp of homology both 5’ and 3’ of the *CDA2* open reading frame. *SAT1* confers resistance to the drug, nourseothricin (NAT). By selecting for NAT resistant transformants, diploids homozygous for *cda2∆* were obtained. Chlamydospore formation was induced in the *cda2∆/cda2∆* diploid on glycerol medium and examined by Eosin Y staining. No Eosin Y staining was observed, confirming the presence of chitosan in the chlamydospore wall (Figure 2B).

To test whether the chlamydospore wall of *C. dubliniensis* also contains dityrosine, chlamydospores were analyzed by fluorescence microscopy using a filter cube optimized for dityrosine (48). Unlike earlier reports in *C. albicans*, no fluorescence was seen specifically in the cell wall, though fluorescence was visible throughout the cytoplasm that was brighter than background fluorescence in the hyphal cells (Figure 2C). This fluorescence is not due to dityrosine, however, as deletion of the *C. dubliniensis DIT1* or *DIT2* genes (which are required for making dityrosine in budding yeast) also exhibited the cytoplasmic fluorescence (Figure 2C). Therefore, a common feature in chalmydospores from *C. dubliniensis* and *C. albicans* and the ascospores from budding yeast is the presence of a chitosan layer in the cell wall.

### A chitosan synthesis pathway is conserved in *C. dubliniensis*

*S. cerevisiae* encodes three different chitin synthases, but chitin synthase 3 (*CHS3*) is specifically used in the synthesis of the chitosan layer of the spore wall (28). *C. dubliniensis*, encodes four different predicted chitin synthases and the ORF Cd36_12160 encodes the ortholog of *S. cerevisiae CHS3* (49, 50). To examine if the use of the Chs3 ortholog for chitosan synthesis is conserved, a *C. dubliniensis chs3*∆/*chs3*∆ mutant was constructed and chlamydospores were stained with Eosin Y. Interestingly, as for the *cda2*∆/*cda2*∆ mutant, greatly reduced fluorescence signal from the Eosin Y staining was seen in the *chs3*∆/*chs3*∆ chlamydospore wall (Figure 3A). In *S. cerevisiae*, deletion of *CDA1* and *CDA2* leads to the accumulation of chitin in the ascospore wall that stains brightly with the dye Calcofluor White (CFW) (29). By contrast, in *C. dubliniensis*, deletion of *CDA2* or *CHS3* does not result in an increase in uniform staining around the cell wall as it does in ascospores (Figure 3A). Rather, CFW predominantly stains the septa consistent with earlier reports in *C. albicans* that chitin at the septum is deposited by chitin synthase 2 (51)(Figure 3A). In sum, these results indicate that Chs3 and Cda2, the same enzymes that generate chitosan in ascopores, collaborate to generate chitosan in the chlamydospore wall,.

**Figure 3.**
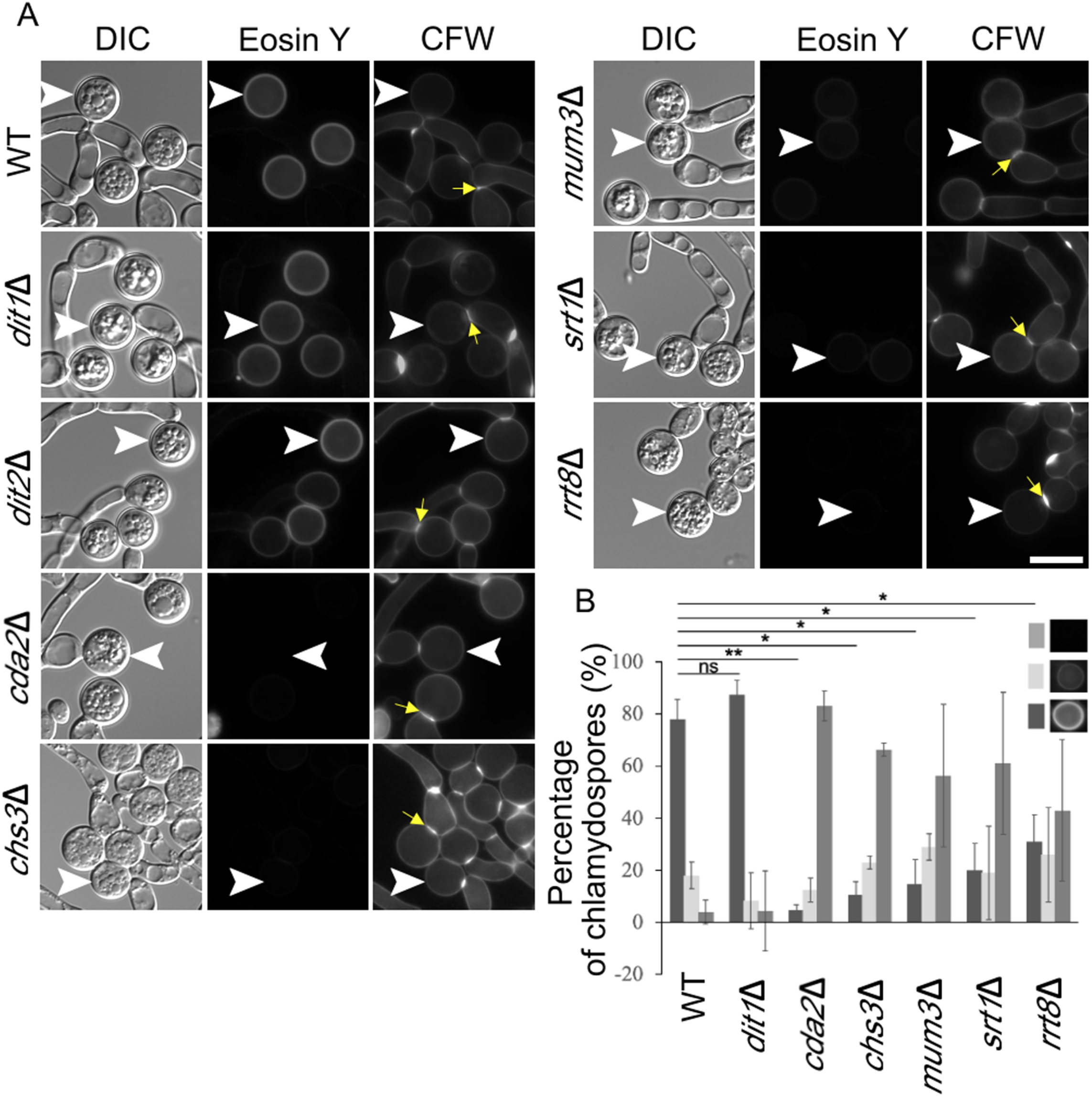
Effect of mutations in *C. dubliniensis* orthologs of *S. cerevisiae* spore wall genes on the chlamydospore wall. (A) Cells of strains of the indicated genotype were grown on SGlycerol medium and then stained with both Eosin Y to label chitosan and Calcofluor White (CFW) to label chitin or chitosan. White arrowheads indicate examples of chlamydospores visible in the images. Yellow arrowheads indicate examples of CFW-stained septa. Scale bar = 10µm. (B) The intensity of the Eosin Y fluorescence was categorized as bright, dim, or no fluorescence for each chlamydospore and the number of chlamydospores in each category for each strain was quantified. For each strain, the value represents the average for one hundred chlamydospores in each of three independent experiments. Error bars indicate one standard deviation. One asterisk (^*^) indicates significant difference at *p<0*.*05*; two asterisks (^**^) indicates significant difference at *p<0*.*0005* Student’s *t*-test.

*C. dubliniensis* encodes uncharacterized orthologs for several of genes required for making ascospore outer cell walls. If the process of chitosan assembly in the wall is conserved, then these same genes may function in chitosan deposition into the chlamydospore wall as well. In particular, we focused on the orthologs of *S. cerevisiae MUM3* (Cd36_82000), *SRT1* (Cd36_11510), and *RRT8* (Cd36_33980). Homozygous deletions for all three of the *C. dubliniensis* genes were constructed and chlamydospores of the mutant strains were examined by Eosin Y and CFW staining. Relative to wild type, the intensity of the Eosin Y fluorescence was reduced in all of the mutant strains, while the fluorescence from CFW staining was unaltered (Figure 3A). These results are similar to the effects of *chs3*∆ and *cda2*∆ and suggest that these genes are important for chitosan formation in *C. dubliniensis*.

To more carefully assess the effect of the mutants, the fluorescence intensity of the Eosin Y staining of individual chlamydospores was categorized as bright, reduced, or absent and the number of chlamydospores in each category was scored for each strain (Figure 3B). The *cda2*∆/*cda2*∆ and *chs3*∆/*chs3*∆ mutant strains displayed a sharp reduction in the fraction of chlamydospores with bright fluorescence intensity and a corresponding increase in chlamydospores displaying no Eosin Y fluorescence. As expected, mutation of *DIT1* or *DIT2* had no obvious effect on Eosin Y staining. By contrast, the *mum3*∆/*mum3*∆, *srt1*∆/*srt1*∆ and *rrt8*∆/*rrt8*∆ diploids all showed phenotypes similar to *chs3*∆ and *cda2*∆ with a significant, though not quite as strong, reduction in brightly staining spores and an increase in unstained spores (Figure 3B).

To confirm that the loss of Eosin Y staining was due to the deletion alleles and not an off target effect from CRISPR/Cas9, the ability of the wild-type gene to complement each mutant was tested. Each wild-type gene was cloned into the integrating plasmid CIp10-SAT, which can be targeted to integrate into the *RPS1* locus (52). This vector uses the same *SAT1* selectable marker that was used to make the deletion alleles. Therefore, prior to transformation with the plasmids, the *SAT1* genes at both copies of each deletion had to be removed. This removal was possible because the knockout cassette included not only the *SAT1* gene but also a maltose-inducible *FLP* recombinase gene, both of which are flanked by flippase recognition target (FRT) sites (46). Induction of the *FLP* recombinase on maltose medium results in recombination between the FRT sites, thereby deleting the *SAT1* and *FLP* genes. Recombinants that lost both copies of *SAT1* were detected by identification of NAT-sensitive colonies. Introduction of *CHS3, CDA2, MUM3*, or *SRT1* into the corresponding knockout strains restored Eosin Y staining to the chlamydospores, confirming that the phenotypes are caused by loss of the specific gene function (Figure 4). We were unable to do the complementation experiment for *rrt8*∆ as the deletion strain failed to grow on the maltose medium used to induce the *FLP* recombinase. Whether the maltose phenotype is a property of the *RRT8* knockout or due to some other change in the strain is unknown. In sum, these results demonstrate that *CHS3, CDA2, MUM3, SRT1*, and probably *RRT8* all contribute to formation of a chitosan component of the chlamydospore wall, suggesting that they constitute a conserved machinery mediating chitosan synthesis for incorporation into yeast cell walls.

**Figure 4.**
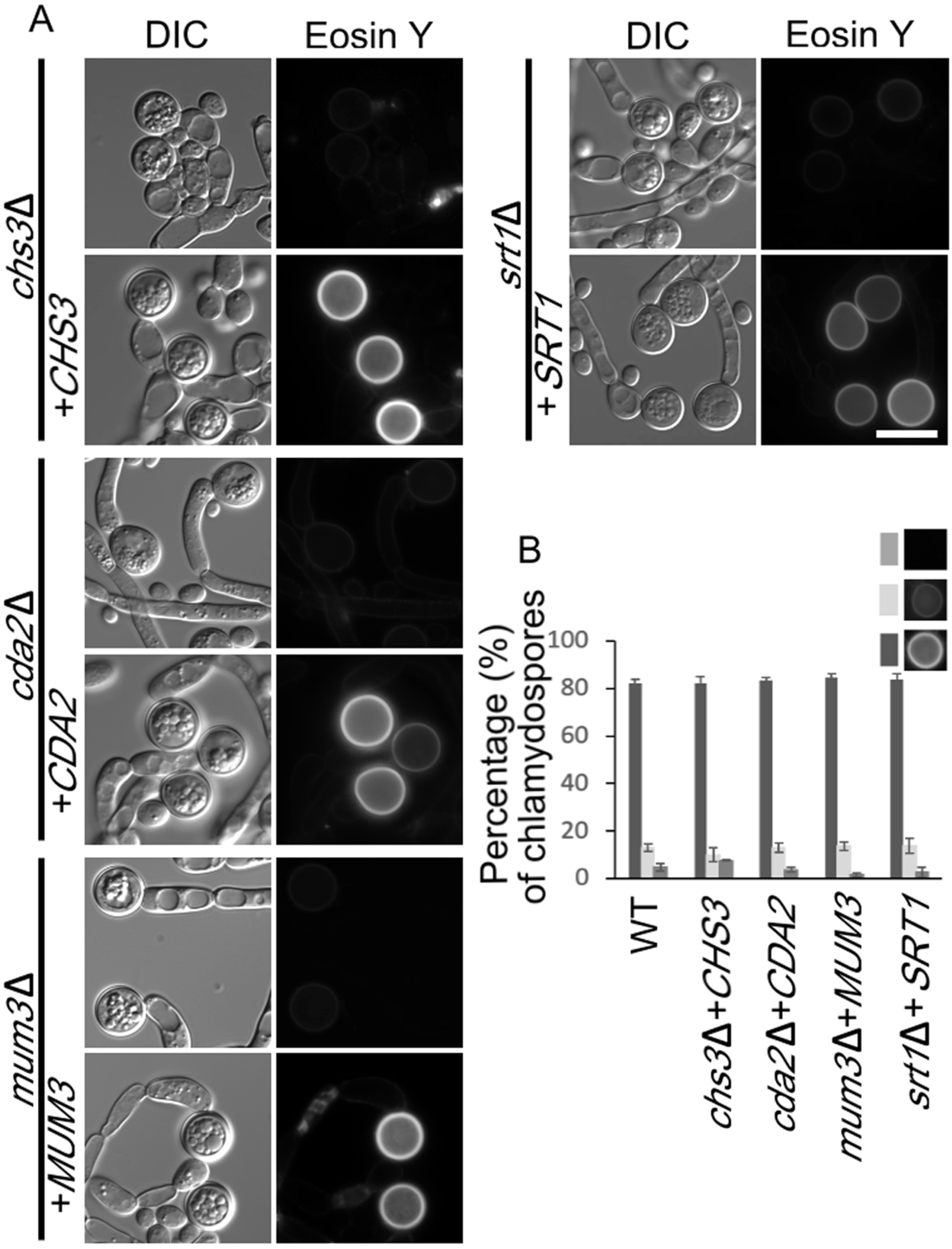
Complementation of the chitosan defect by the wild-type alleles. (A) A wild type copy of *CHS3, CDA2, MUM3*, or *SRT1* gene, respectively, was integrated into the corresponding deletion mutant (strains BEM15-18). Cells were grown on SGlycerol medium and Eosin Y staining of chlamydospores with our without reintroduction of the wild-type allele was examined. DIC = differential interference contrast. Scale bar = 10µm. (B) Rescue of Eosin Y staining by the wild-type alleles was quantified as in Figure 3.

### Ultrastructural analysis identifies a chitosan layer in the chlamydospore wall

The fluorescence images from the Eosin Y staining suggest that chitosan is missing or reduced in the chlamydospore wall of various mutants. Previous ultrastructural studies have revealed that the chlamydospore wall of *C. albicans* is distinct from the hyphal wall in having a darkly staining inner layer of unidentified material underneath what appear to be beta-glucan and mannan layers (42, 53). To examine the ultrastructure of the *C. dubliniensis* chlamydospore wall, cells were stained using osmium and thiocarbohydrazide and examined by electron microscopy (31). Similar to previous reports, the cell wall of wild-type chlamydospores displayed a layer of darkly staining material close to the plasma membrane with outer, lighter layers resembling the walls of the adjacent hyphal cells (Figure 5A, B).

**Figure 5.**
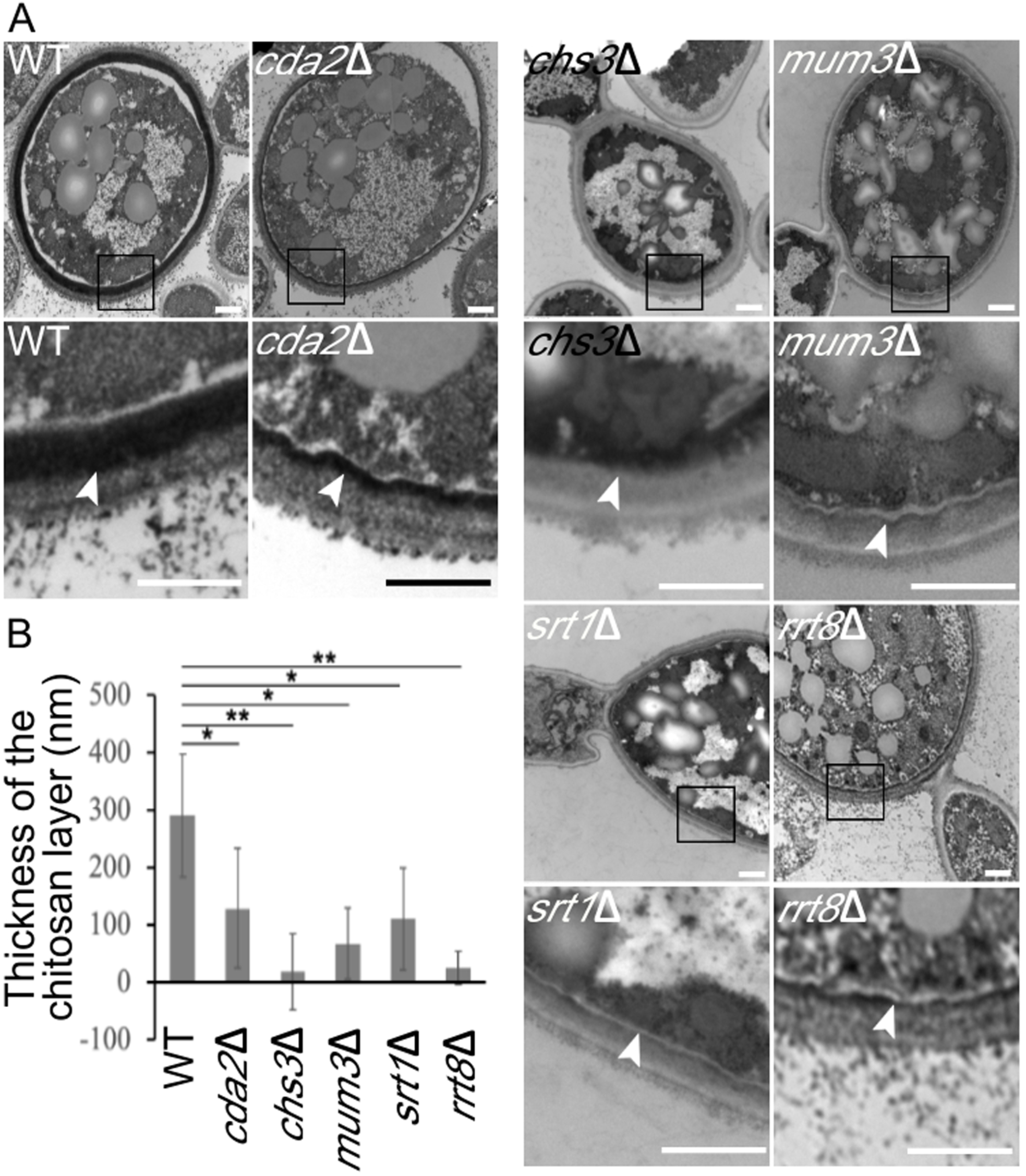
Electron microscopy of the chamydospore wall of *C. dubliniensis*. (A) Chlamydospores were induced and cells of different strains were stained with osmium-thiocarbohydrazide: WT (CD1465), *cda2*Δ (BEM7), *chs3*Δ (BEM8), *mum3*Δ (BEM11), *srt1*Δ (BEM12), *rrt8*Δ (BEM13). For each strain, a pair of images is shown. The lower image is a higher magnification of the boxed region in upper image. Arrowheads indicate the inner cell wall layer. (B) Quantification of the thickness of the chitosan layer in each strain. Data represented are the means of measurements from 20 chlamydospores. The thickness of the chitosan layer was measured at 5 different positions on each chlamydospore. Error bars indicate one standard deviation. One asterisk (^*^) indicates significant difference at *p<0*.*00005*; two asterisks (^**^) in indicates *p<5e-10*, Students *t*-test. Scale bar = 500 nm

Given that the chitosan-containing outer ascospore wall of *S. cerevisiae* also stains darkly under these conditions (31), this inner, electron dense material in the chlamydospore wall may be chitosan. Consistent with this possibility and with the Eosin Y fluorescence results, this inner layer was dramatically reduced in both the *chs3*∆/*chs3*∆ and *cda2*∆/*cda2*∆ strains (Figure 5A). Thus, as in the ascospore wall, chitosan in the chlamydospore wall forms a discrete layer. Again, consistent with the Eosin Y fluorescence results, the chitosan layer appeared reduced or absent in chlamydospores of the *mum3*∆, *srt1*∆, and *rrt8*∆ mutants as well (Figure 5A).

The reduction in the chitosan layer visible in the electron micrographs was somewhat variable between chlamydospores in individual strains. Therefore, to measure the effect of the mutants, the thickness of the chitosan layer in the micrographs was measured as an indicator of the amount of chitosan deposited. In each strain, the thickness of the chitosan layer was measured at five locations in twenty different chlamydospores (Figure 5B). All of the mutants displayed significantly reduced chitosan layers, with *chs3*∆ displaying the strongest phenotype. In sum, the ultrastructural analysis confirms that chitosan is present in a discrete layer of the chlamydospore wall and a conserved set of genes is required for proper formation of this layer.

### *C. dubliniensis* Rrt8, Mum3, and Srt1 are all localized on lipid droplets

In *S. cerevisiae*, the Srt1 and Rrt8/Lds1/Lds2 proteins are localized to lipid-droplets, and lipid droplets are associated with the forming spore wall, suggesting some connection between lipid droplets and the assembly of the outer spore wall layers (34-36). *C. albicans* chlamydospores are reported to be rich in neutral lipids and lipid droplets based on both biochemical fractionation and staining with a lipid droplet dye (45, 54). To examine lipid droplets in *C. dubliniensis* chlamydospores, the cells were stained with the lipid droplet dye monodansylpentane (MDH) (55). This treatment revealed a very high density of lipid droplets within the chlamydospore as compared to *C. dubliniensis* cells growing in yeast phase that was not changed in any of the mutant strains (Figure 6).

**Figure 6.**
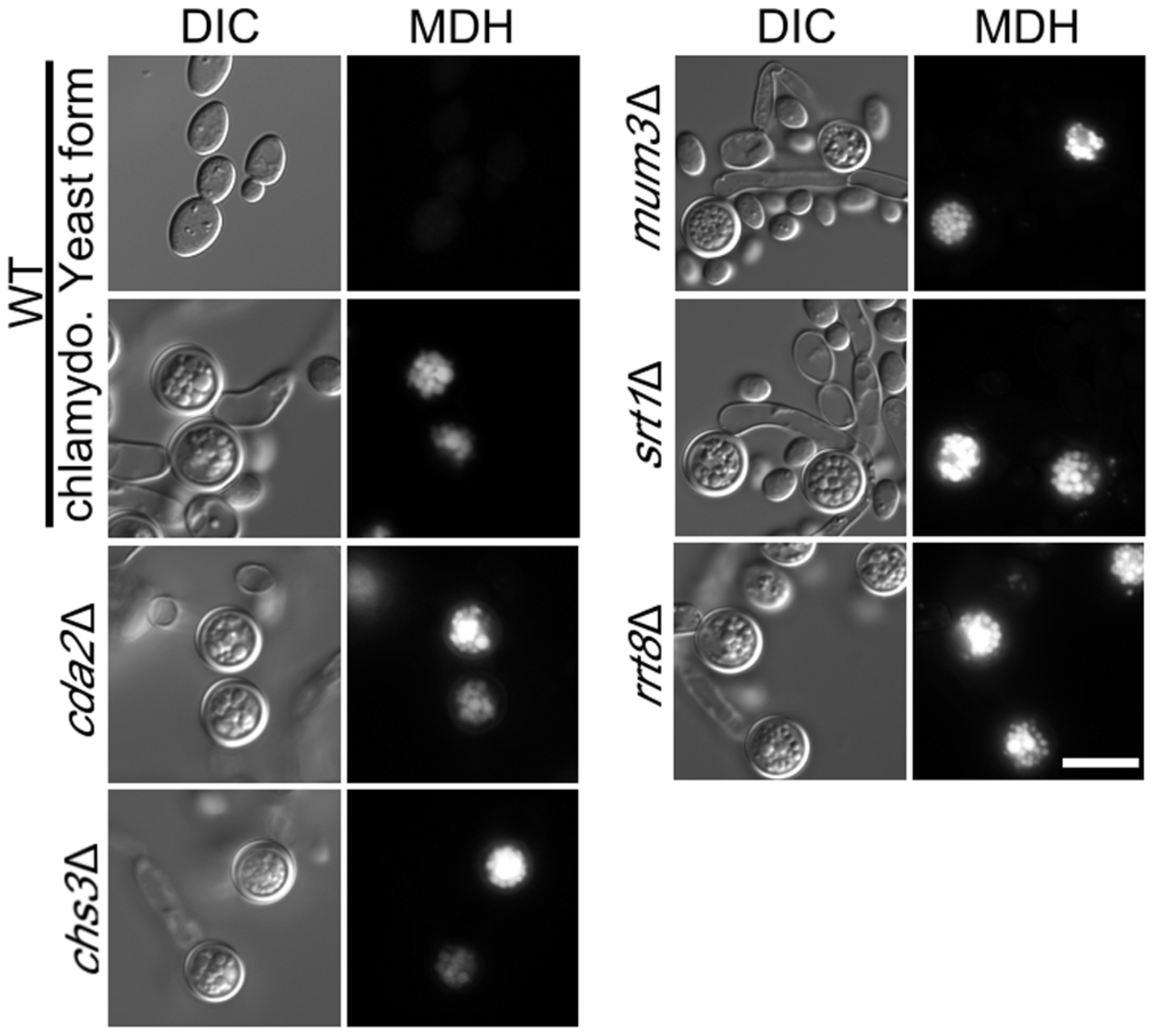
Lipid droplets in chlamydospores. WT cells (CD1465) growing on SGlucose or SGlycerol media or the indicated mutant strains (BEM7, 8, 11, 13, 14) grown on SGlycerol were stained with MDH to label lipid droplets and visualized using a BFP filter. Scale bar = 10µm

The abundance of lipid droplets in the chlamydospore and the connection of the *S. cerevisiae* proteins to lipid droplets led us to examine the localization of the different *C. dubliniensis* proteins. Each gene, under the control of its native promoter, was fused at its 3’ end to a gene encoding a *Candida* codon-optimized red fluorescent protein (yEmRFP) (56). Plasmids containing the fusion genes were then integrated at the *RPS1* locus in the appropriate deletion strains (except for *rrt8∆* where we were unable to eliminate the *SAT1* gene from the deletion, so a wild-type strain was used). *C. dubliniensis* cells carrying the different *yEmRFP* fusions were then grown under chlamydospore-inducing conditions, stained with MDH to detect lipid droplets, and examined by fluorescence microscopy. For the *MUM3, SRT1*, and *RRT8* fusions, red fluorescence co-localized with the lipid droplet marker in the chlamydospores (Figure 7A). Red fluorescence at the the cell periphery was also visible in the wild-type strain carrying no yEmRFP and so is background fluorescence visible due to the longer exposures necessary to visualize the yEmRFP fusions. Importantly, no background fluorescence was seen at the lipid droplets.

**Figure 7.**
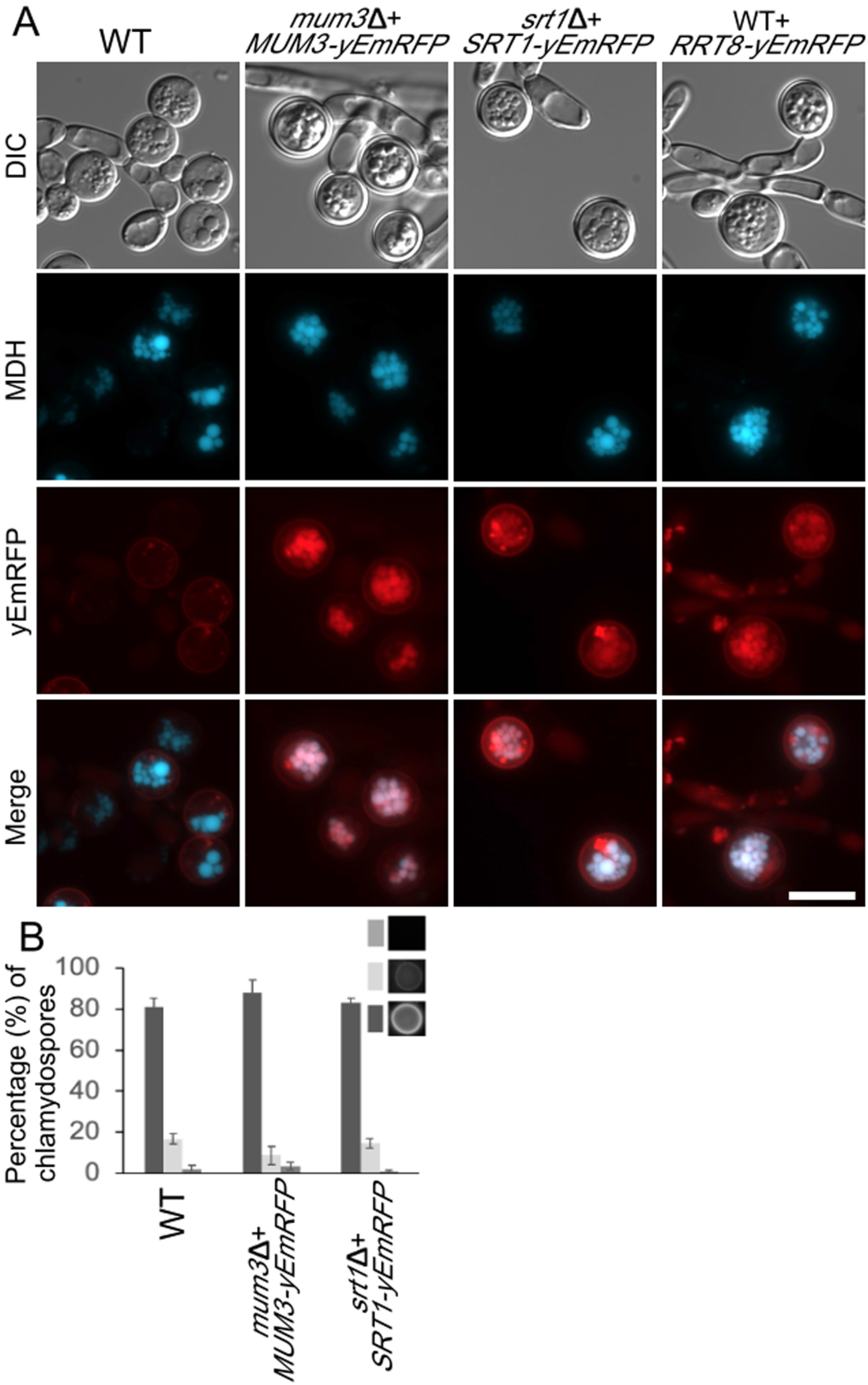
Localization of Cda2, Mum3, Rrt8 and Srt1 in chlamydospores. WT (Cd1456) cells expressing no RFP fusion or strains expressing different *MUM3*-, *SRT1*, or *RRT8*-*yEmRFP* fusions (BEM20, 21, 22) were grown on SGlycerol medium, stained with MDH and visualized through both BFP and RFP filters. Scale bar = 10µm ((B) Eosin Y staining of chlamydospores in WT (CD1465) *mum3*∆ *MUM3-yEMRFP* (BEM20) and *srt1*∆ *SRT1-yEmRFP* (BEM22) strains was quantified as in Figure 3.

To confirm that the fusion proteins are functional, the appropriate deletion strains carrying *MUM3::yEmRFP* or *SRT1::yEmRFP* were examined for the ability of the fusion to rescue the mutant phenotype by staining of chlamydospores with Eosin Y (Figure 7B). Both fusions restored bright Eosin Y staining indicating that the lipid droplet-localized fusion proteins are functional. The localization of all three proteins suggests that lipid droplets promote chitosan layer formation in *C. dubliniensis*.

## Discussion

We report that *C. dubliniensis* efficiently forms chlamydospores when incubated on synthetic medium containing different non-fermentable carbon sources. While the molecular signals that trigger chlamydosporulation are complex (40), nutritional signals are known to be involved and induction by changing carbon sources suggests that central carbon metabolism may play a role. Whether this induction mechanism is unique to *C. dubliniensis* remains to be seen, since *C. albicans* was not induced to form chlamydospores under these conditions. Previous studies reported that *C. dubliniensis* can form chlamydospores in Staib medium (a seed extract) (57). Wild-type *C. albicans* does not form chlamydospores efficiently under these conditions, but deletion of the *C. albicans NRG1* gene leads to chlamydosporulation in Staib medium similar to *C. dubliniensis* (41). The signals triggering chlamydosporulation may be different in SGlyerol and Staib medium, however, as no chlamydospores were seen on SGlycerol when a *C. albicans nrg1* mutant was used (L.D.B., unpublished observation).

To create mutant strains in *C. dubliniensis*, we utilized a transient CRISPR-Cas9 system originally developed for *C. albicans* (46). Combining this transient system with the recyclable *SAT1-FLP* cassette allowed us to do multi-step strain constructions directly in clinical isolates without the need for auxotrophic markers, greatly accelerating our analysis. That this system works well in both *C. dubliniensis* and *C. albicans* suggests that it will be useful for other *Candida* species as well.

In contrast to a previous report about the chlamydospore wall of *C. albicans* (44), we see no evidence that the *C. dubliniensis* chlamydospore wall contains dityrosine. In the earlier study, dityrosine fluorescence of the *C. albicans* chlamydospore wall was observed in wild-type cells and deletion of the *CYP56*/*DIT2* gene abolished chlamydospore formation (44). By contrast, the *C. dubliniensis dit2*∆ mutant formed abundant chlamydospores (Figure 2). It is possible these results represent a difference between *C. albicans* and *C. dubliniensis*, though because the *C. albicans dit2*∆ mutant did not form chlamydospores, it is not clear whether the fluorescence observed in the wild-type *C. albicans* chlamydospores was dityrosine or fluorescence from some other molecule.

We show here that chitosan is a major constituent of the previously described dark, inner layer of the *Candida* chlamydospore wall. Chitosan also forms a discrete layer in the *S. cerevisiae* ascospore wall, however, the position of the chitosan layer with respect to other cell wall components is distinct in the two cell walls (Figure 8). In the ascospore, the chitosan is located towards the outside of the structure while in the chlamydospore it is on the interior of the wall. In both cases, however, the chitosan is localized adjacent to the beta-glucan components of the wall, suggesting that the presence of the beta-glucan may also be important for organizing the chitosan into a distinct layer.

**Figure 8.**
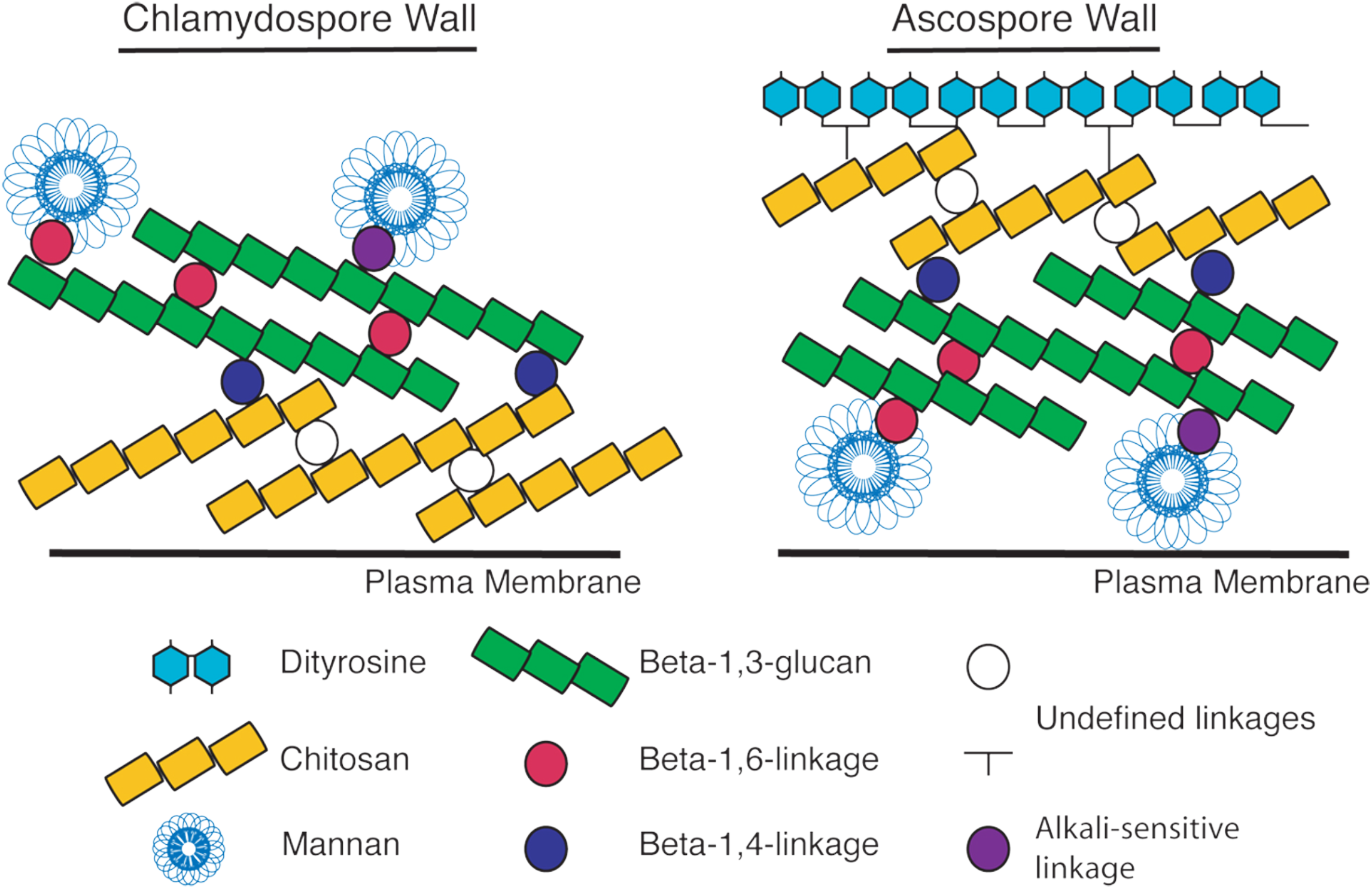
Model for organization of the *C. dubliniensis* chlamydospore and *S. cerevisiae* ascospore walls. The organization of the different layers of the walls are shown with respect to the cell plasma membrane. The linkages between components are based on the known linkages in the vegetative cell wall of *S. cerevisiae* (25). The nature of the crosslinks within and between the chitosan and dityrosine layers is unknown.

Our results reveal a conserved machinery required for chitosan layer synthesis. Multiple chitosan synthases are present in both *S. cerevisiae* and *C. dubliniensis* and yet in both yeasts, *CHS3* is uniquely required for synthesis of the chitosan layer of the ascopore and chlamydospore cell walls. Whether this reflects a specific association of this chitin synthase with the chitin deacetylase protein or with some other aspect of Chs3 activity remains to be determined. For example, the Chs3 enzyme might synthesize chitin strands of a chain length or organization that is more amenable to deacetylation. Indeed, *C. albicans* Chs3 has been reported to synthesize shorter chitin fibrils than Chs8 (50).

The lipid-droplet localized proteins Srt1, Rrt8 and Mum3 are required for proper chitosan layer formation in both yeasts. As these proteins are localized on cytosolic lipid droplets, their effects on chitosan assembly must be somewhat indirect. *MUM3* and *SRT1* encode predicted lipid-synthesizing enzymes. The Mum3 protein is homologous to O-acyltransferase enzymes and Srt1 is a subunit of a cis-prenyltransferase responsible for synthesizing a lipid-droplet localized pool of polyprenols (31, 34). In earlier work, we proposed a model in which Srt1-generated long chain polyprenols in the lipid droplet that are transferred to the plasma membrane to enhance Chs3 activity (34). It is possible that a similar mechanism occurs during chlamydospore formation. An alternative possibility was recently suggested by nuclear magnetic resonance (NMR) studies of chitosan-containing cell wall preparations from both *S. cerevisiae* and *Cryptococcus neoformans* that revealed neutral lipids are directly incorporated into the cell wall (16, 58). Thus, the Rrt8, Srt1, and Mum3 proteins may be involved in the synthesis of some lipid component that is then transferred from the lipid droplet to play a structural role during chitosan layer assembly. Further biochemical work will be necessary to clarify how these proteins and their lipid products contribute to formation of this cell wall structure.

NMR studies suggest that there is a common architecture for chitosan-containing elements in the fungal cell wall from ascomycetes to basidiomycetes (16). Orthologs of the genes described here that underlie formation of chitosan cell wall layers in *Candida* and *Saccharomyces* can be found throughout the fungi. Thus, the similar architecture may reflect a broadly conserved genetic network regulating the synthesis of chitosan-containing cell wall structures in fungi. Given the importance of chitosan to virulence of some pathogenic fungi, the genes described here may be useful potential targets for antifungal therapies (13).

## Materials and methods

### Strain and growth conditions

Strain used are listed in Table 1. *C. dubliniensis* strain Cd1465 is derived from a clinical specimen isolated from a patient sample at the Stony Brook hospital. This strain was routinely cultured at 30ºC on YPD medium (2% Bacto peptone, 2% dextrose, 1% yeast extract and 2% agar). *C. dubliniensis* transformants were selected on YPD_NAT (2% Bacto peptone, 2% dextrose, 1% yeast extract, 2% agar and 400μg/ml nourseothricin [Werner BioAgents]) for nourseothricin-resistant isolates. Synthetic glycerol (SGlycerol) solid medium 1.7% yeast nitrogen base without amino acids, 2% agar and 0.1M glycerol) was used to induce chlamydospores, as described below.

**Table 1.** Strains used in this study

| Strain | Genotype | Reference |
| --- | --- | --- |
| Cd1465 | Wild-type | This study |
| Cd1466 | Wild-type | This study |
| Cd1467 | Wild-type | This study |
| BEM7 | Cd1465, plus *cda2Δ::FRT-SAT1::FLIP-FRT /cda2Δ:: FRT-SAT1::FLIP-FRT* | This study |
| BEM8 | Cd1465, plus *chs3Δ:: FRT-SAT1::FLIP-FRT /chs3Δ:: FRT-SAT1::FLIP-FRT* | This study |
| BEM9 | Cd1465, plus *dit1Δ:: FRT-SAT1::FLIP-FRT /dit1Δ:: FRT-SAT1::FLIP-FRT* | This study |
| BEM10 | Cd1465, plus *dit2Δ:: FRT-SAT1::FLIP-FRT /dit2Δ:: FRT-SAT1::FLIP-FRT* | This study |
| BEM11 | Cd1465, plus *mum3Δ:: FRT-SAT1::FLIP-FRT /mum3Δ::FRT-SAT1::FLIP-FRT* | This study |
| BEM13 | Cd1465, plus *rrt8Δ:: FRT-SAT1::FLIP-FRT /rrt8Δ:: FRT-SAT1::FLIP-FRT* | This study |
| BEM14 | Cd1465, plus *srt1Δ:: FRT-SAT1::FLIP-FRT /srt1Δ:: FRT-SAT1::FLIP-FRT* | This study |
| BEM15 | Cd1465, plus *cda2Δ::FRT/cda2Δ::FRT RPS1::P<sub>CDA2</sub>CDA2-Clp10-SAT1/RPS1* | This study |
| BEM16 | Cd1465, plus *chs3Δ::FRT/chs3Δ::FRT RPS1::P<sub>CHS3</sub>CHS3-Clp10-SAT1/RPS1* | This study |
| BEM17 | Cd1465, plus *mum3Δ::FRT/mum3Δ::FRT RPS1::P<sub>MUM3</sub>MUM3-Clp10-SAT1/RPS1* | This study |
| BEM18 | Cd1465, plus *srt1Δ::FRT/srt1Δ::FRT RPS1::P<sub>SRT1</sub>SRT1-Clp10-SAT1/RPS1* | This study |
| BEM19 | Cd1465, plus *cda2Δ::FRT/cda2Δ::FRT RPS1::P<sub>CDA2</sub>CDA2-yEmRFP-Clp10-SAT1/RPS1* | This study |
| BEM20 | Cd1465, plus *mum3Δ::FRT/mum3Δ::FRT RPS1::P<sub>MUM3</sub>MUM3-yEmRFP-Clp10-SAT1/RPS1* | This study |
| BEM21 | Cd1465, plus *RPS1::P<sub>RRT8</sub>RRT8-yEmRFP-Clp10-SAT1/RPS1* | This study |
| BEM22 | Cd1465, plus *srt1Δ::FRT/srt1Δ::FRT RPS1::P<sub>SRT1</sub>SRT1-yEmRFP-Clp10-SAT1/RPS1* | This study |

### Induction of chlamydospores

To induce chlamydospores formation, wild-type and mutant strains were inoculated in 5ml YPD liquid and were incubated at 30ºC with shaking at 220rpm for overnight. A suspension of 1 × 10^7^cells/ml was prepared from the overnight culture. The cell suspension was then diluted 100 times and 1ml was spread on a SGlycerol plate. Excess liquid was removed by pipetting and the plates were left to dry at room temperature. All the plates were incubated in 30ºC for 24h. The chlamydospores were collected by adding 500μl of distilled water to the plate and gently scraping the surface of the plate with a glass rod.

### CRISPR-Cas9 mutagenesis in *C. dubliniensis*

To create knockout mutations in *C. dubliniensis*, we adapted a CRISPR/CAS9 system developed for *C. albicans* (59). The pV1093 vector carries both Cas9 and single guide RNA (sgRNA) expression cassettes (59). Guide RNAs targeting specific genes were designed using the CCTop (CRISPR/Cas9 target online) program (60). The *CAS9* gene expression cassette and the sgRNA scaffold were amplified separately from pV1093 using the primers BLD1 and BLD2. The sgRNA scaffold contains the *SNR52* promoter was assembled by the single-joint PCR method (61). Briefly, three-DNA synthesis step was used to generate the sgRNA cassette. The first step consists to amplify by PCR the *SNR52* promoter and sgRNA scaffold using gene-specific flanking primers (Table 2) and internal chimeric primers (BLD3 and BLD4). Twenty complementary bases overlapped and specified the sgRNA of each gene to be knocked out. The second step, both components were fused by primer extension, relying upon annealing of the complementary chimeric primer extensions. The third step consists to amplify the joined product with nested primers (BLD5 and BLD6) to yield the sgRNA cassette.

**Table 2.**
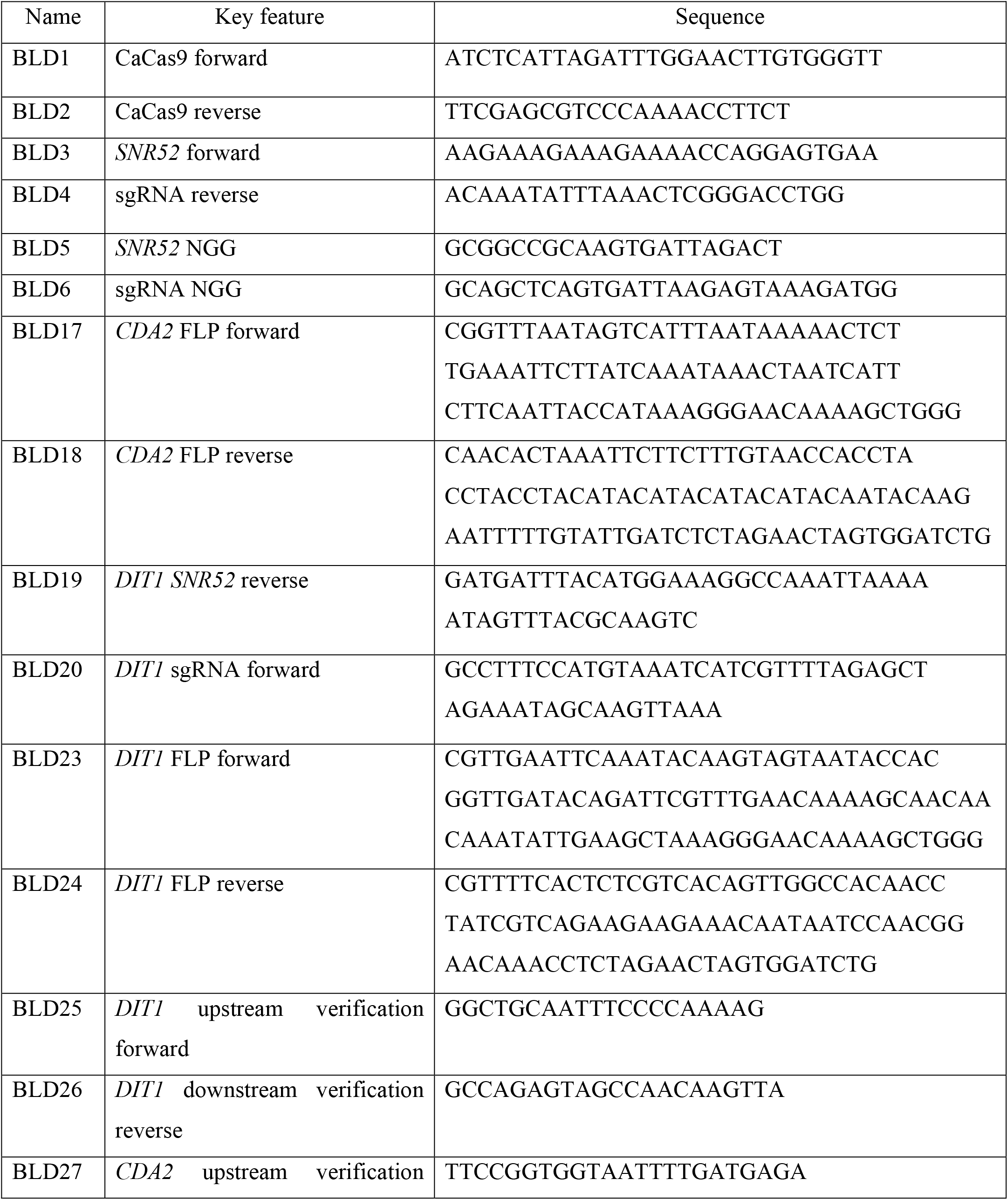
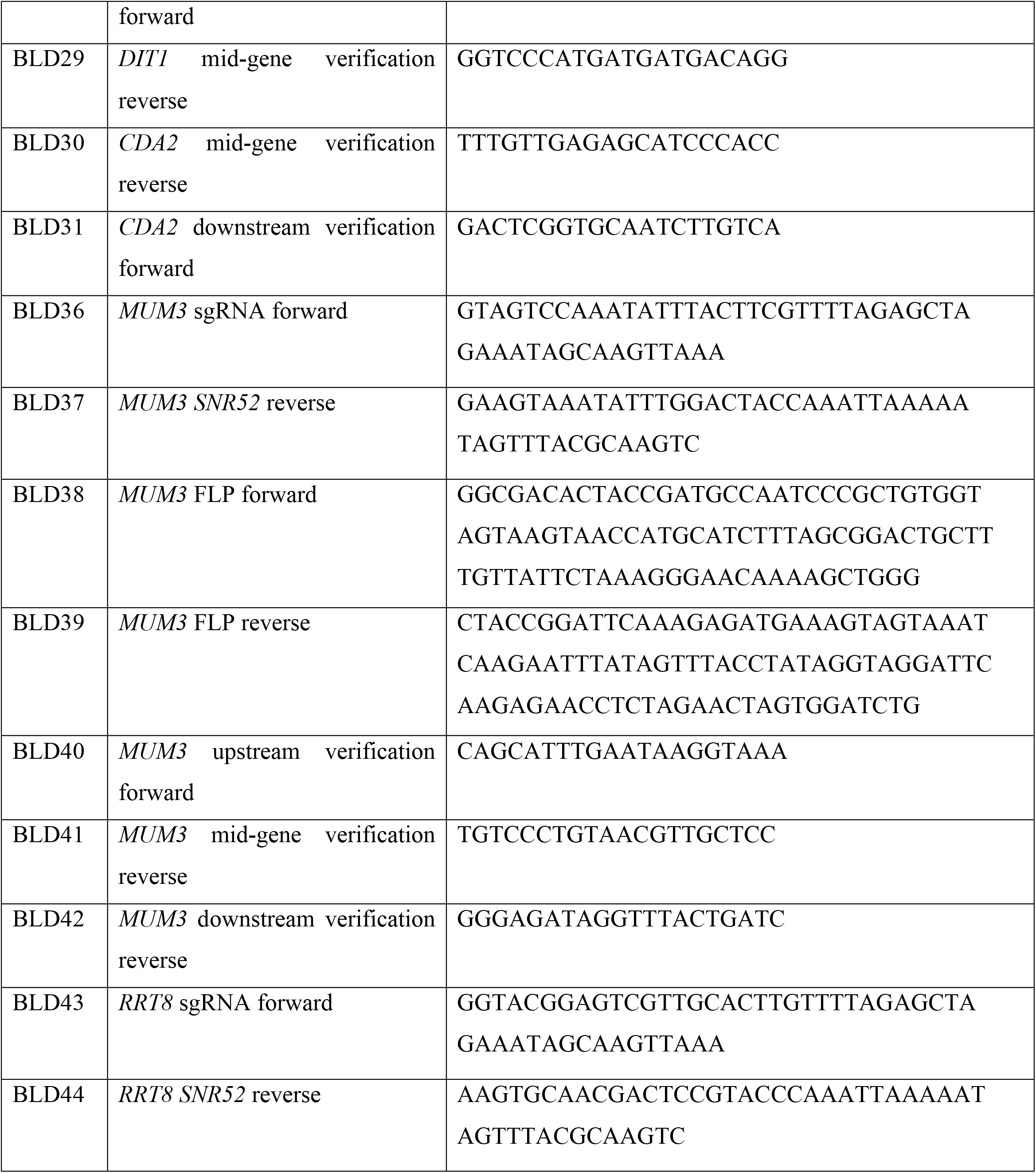
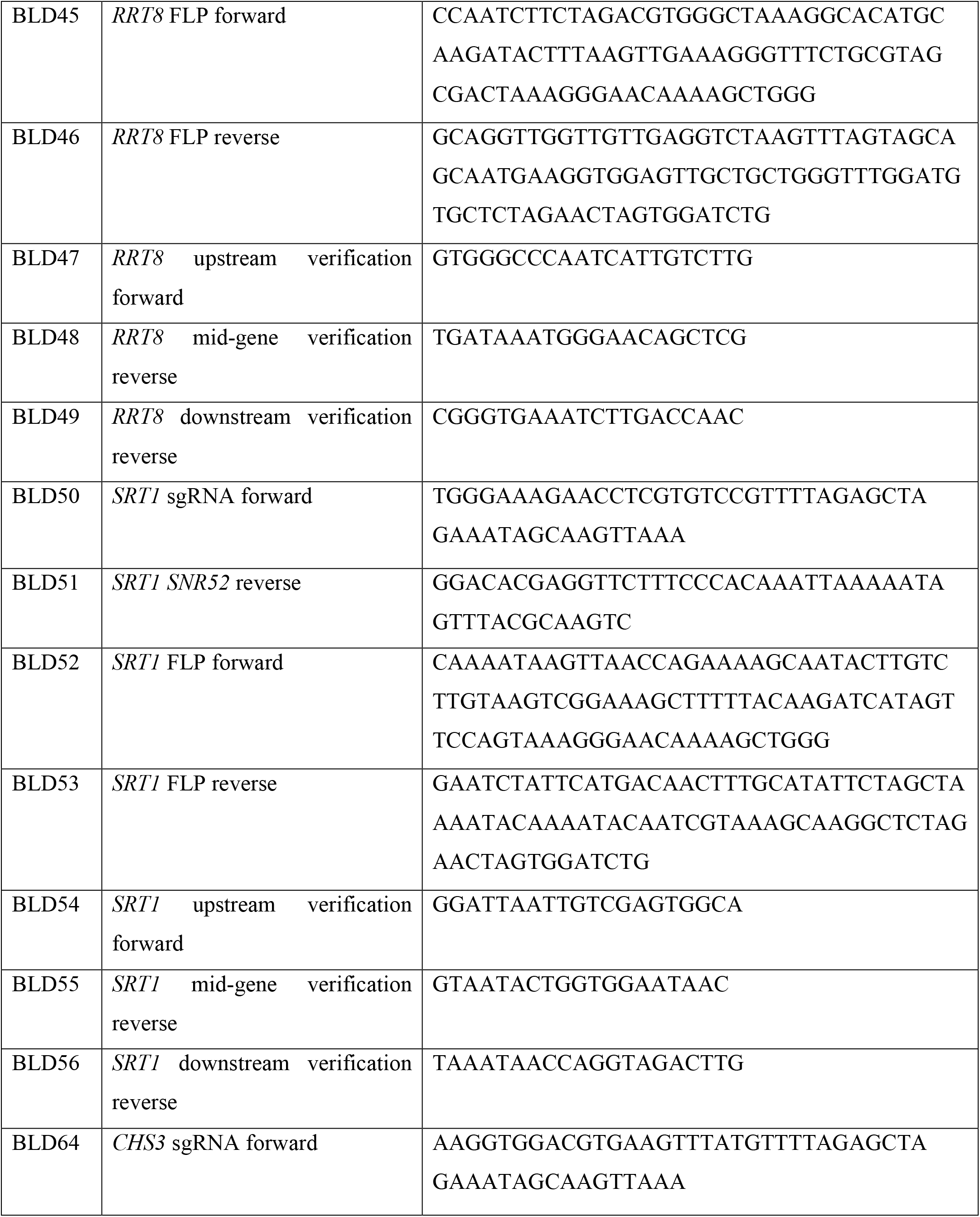
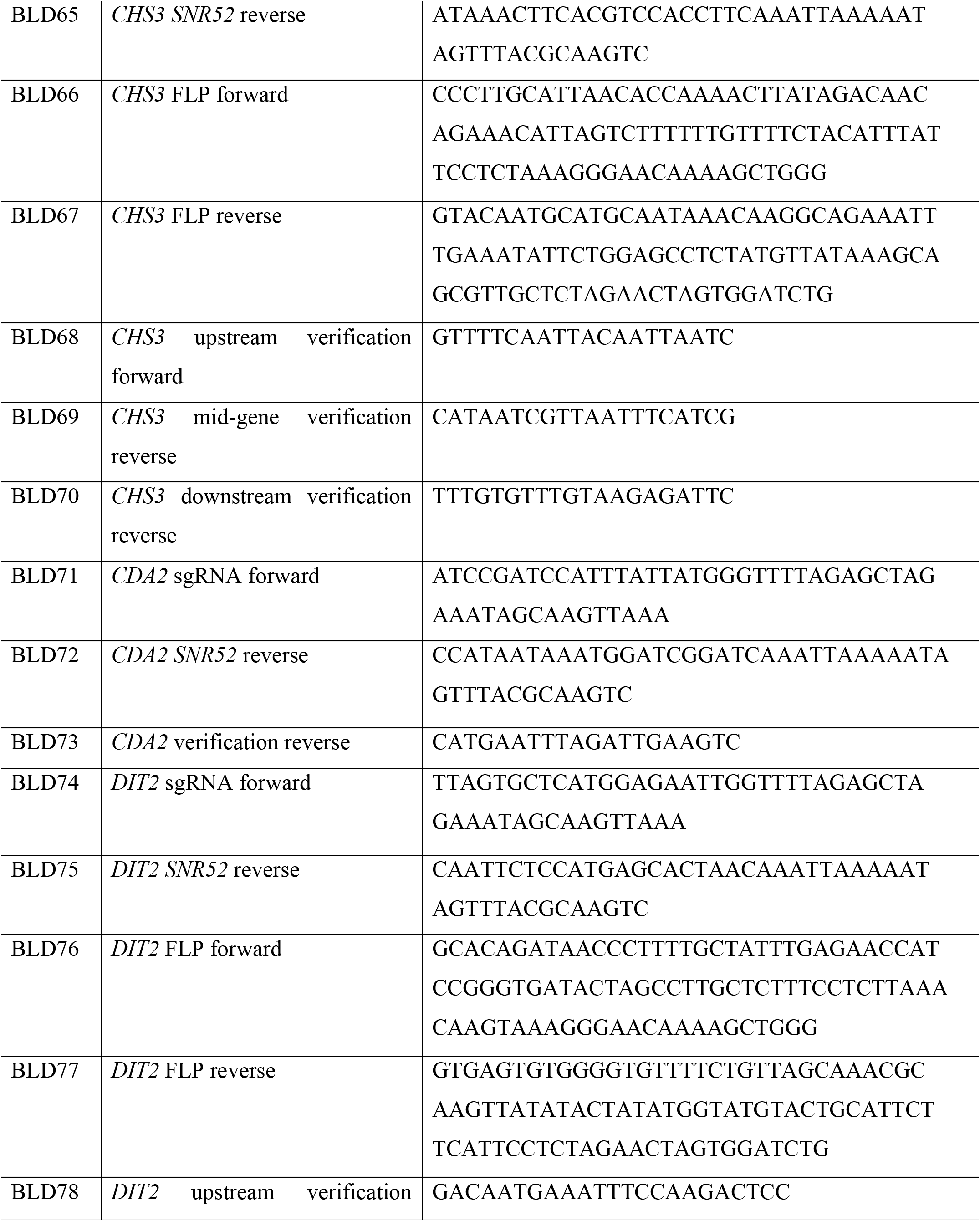
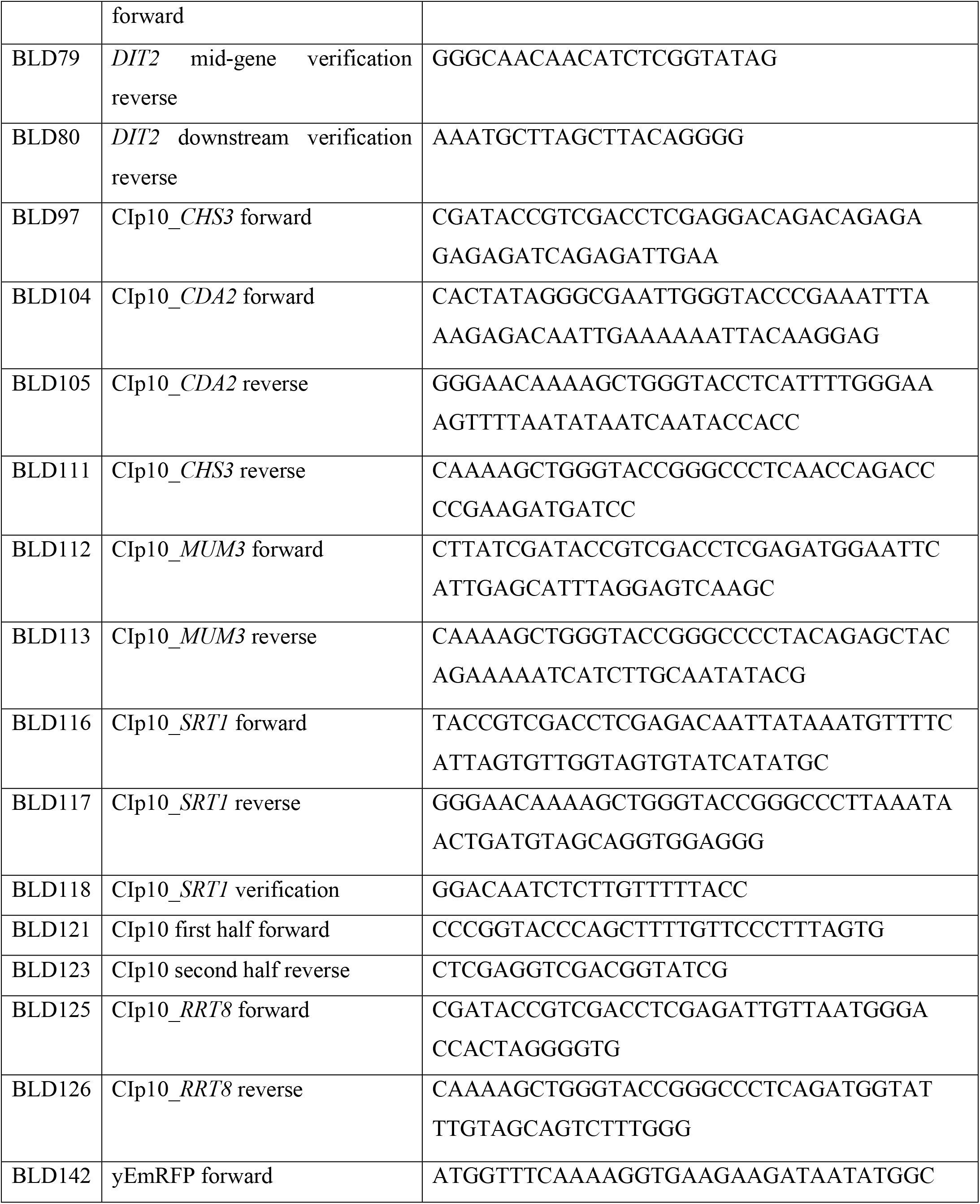
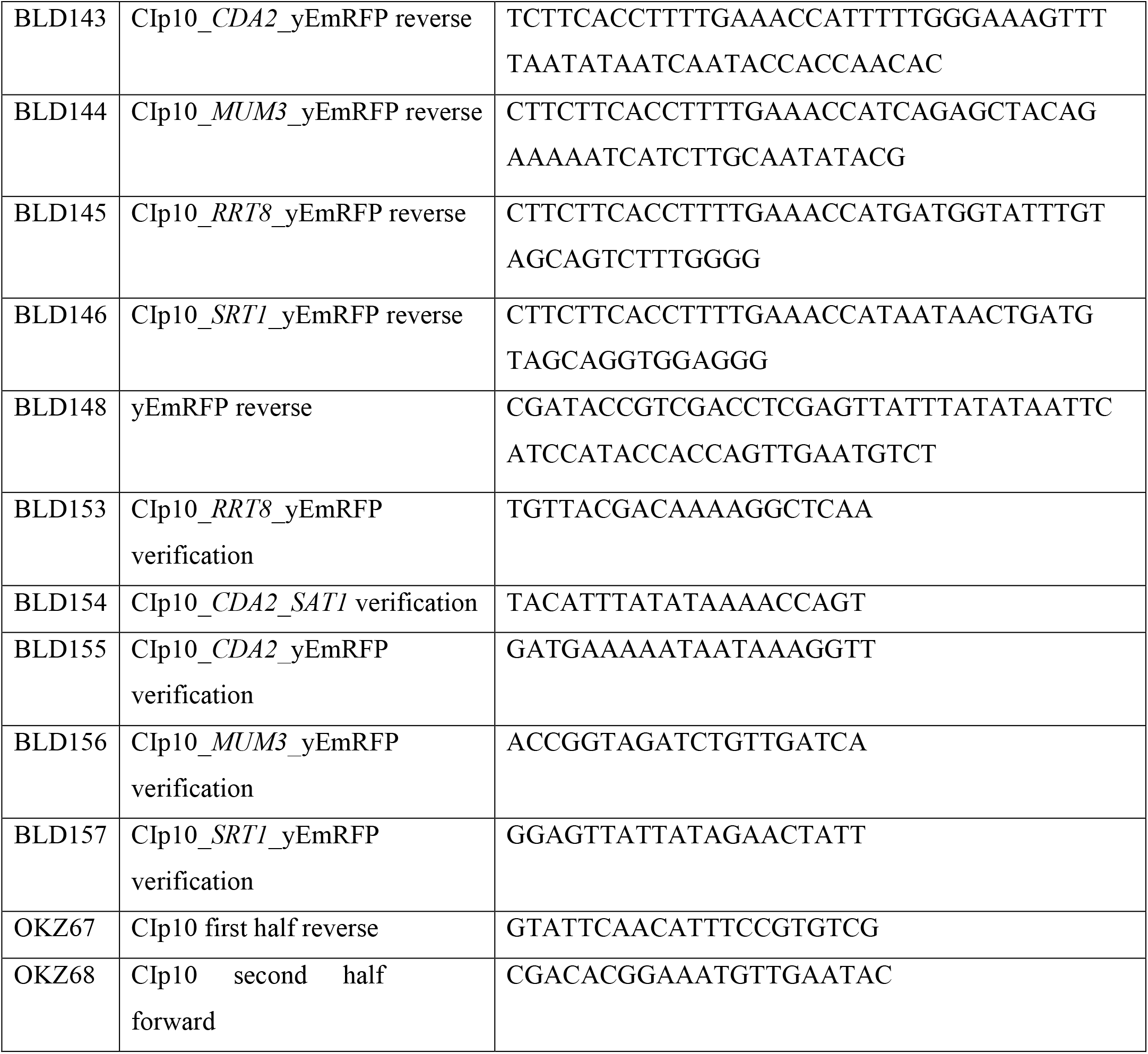
Oligonucleotide primers used in this study

The FLP recombination target sequence target (FRT) and the *SAT1* cassette both encoded in pGR_NAT vector, were flanked by ∼20 bp homology to the 5’and 3’ regions of the gene to be knocked out. This fragment was PCR amplified and used as the gene deletion construct (46). The oligonucleotides used in this study are listed in Table 2. PCR amplification were conducted using Ex *Taq* in accordance with the manufacturer’s instructions (TaKaRa Bio, Inc.).

For the mutagenesis, PCR products for transformation were purified and concentrated with a commercial PCR purification kit (Qiagen, Maryland, USA). The deletion constructs (3μg) were co-transformed with the CdCAS9 cassette (1μg) and the sgRNA cassette (1μg) using the lithium acetate transformation method (62). At least five independent homozygous deletion strains were tested for each mutant.

### Rescue of mutant strains

For each mutant, to confirm that the observed phenotypes were due to the deletion, an integrating plasmid carrying the wild type gene was constructed. CIp10-SAT (a gift of N. Dean) was used as the vector. To construct the complementing plasmids, CIp10 was amplified as two separate fragments by PCR. The first fragment, amplified with BLD121 and OKZ67, contains the *ApaI* site at the one end and part of the Amp locus at the other end. The second fragment, amplified with BLD123 and OKZ68, harbors an overlapping fragment of the Amp locus at one end and an *XhoI* site at the other end. Each gene of interest was amplified by PCR from *C. dubliniensis* genomic DNA with 15bp homologous sequence to the region of CIp10 carrying the *ApaI* or *XhoI* sites at the opposite ends. *CDA2* was amplified with BLD104 and BLD105, *CHS3* with BLD97 and BLD11, *MUM3* with BLD112 and BLD113, and *SRT1* with BLD116 and BLD117. The three fragments were fused by Gibson Assembly (BioLabs) and transformed into *E. coli*. All the plasmids used in this study are listed in Table 3.

**Table 3.** Plasmids used in this study

| Plasmid no. | Name | Key feature | Reference |
| --- | --- | --- | --- |
|  | pNAT | *P<sub>URA3</sub>URA3 SAT1* | (46) |
|  | pV1093 | *CaCas9/SAT1 flipper ENO1* | (59) |
|  | Clp10-SAT | *CaRPS1 SAT1* | N. Dean |
|  | yEpGAP_Cherry | *URA3 yEmRFP* | (56) |
| pLB1 | Clp10_*CDA2* | *CaRPS1 P<sub>CDA2</sub>CDA2 SAT1* | This study |
| pLB2 | Clp10_*CHS3* | *CaRPS1 P<sub>CHS3</sub>CHS3 SAT1* | This study |
| pLB3 | Clp10_*MUM3* | *CaRPS1 P<sub>MUM3</sub>MUM3 SAT1* | This study |
| pLB4 | Clp10_*RRT8* | *CaRPS1 P<sub>RRT8</sub>RRT8 SAT1* | This study |
| pLB5 | Clp10_*SRT1* | *CaRPS1 P<sub>SRT1</sub>SRT1 SAT1* | This study |
| pLB6 | Clp10_*CDA2_yEmRFP* | *CaRPS1 P<sub>CDA2</sub>CDA2 yEmRFP SAT1* | This study |
| pLB7 | Clp10_*MUM3_yEmRFP* | *CaRPS1 P<sub>MUM3</sub>MUM3 yEmRFP SAT1* | This study |
| pLB8 | Clp10_*RRT8_yEmRFP* | *CaRPS1 P<sub>RRT8</sub>RRT8 yEmRFP SAT1* | This study |
| pLB9 | Clp10_*SRT1_yEmRFP* | *CaRPS1 P<sub>SRT1</sub>SRT1 yEmRFP SAT1* | This study |

In order to rescue the mutant strains, we first recycled the selectable marker *SAT1*. To allow the recycling, the mutant strains were plated on YPM [2% Bacto peptone, 2% maltose, 1% yeast extract, 2% agar] to induce expression of the FLP recombinase (47) and then replica-plated to YPD_NAT medium. Colonies that became sensitive to nourseothricin were selected for transformation with the integrating plasmid carrying the corresponding wild type gene. The plasmids were linearized by digestion with NcoI before transformation into the mutant strains by lithium acetate transformation method (62) with modifications. Briefly, fresh overnight cultures (12 h to 16 h) were diluted 1:50 and incubated for ∼ 6 h (optical density at 600 nm [OD_600_] of 5.0. The cells were harvested, washed once with H_2_O and once with 100 mM lithium acetate (LiOAc), and resuspended in 100 µl LiOAc (100 mM). Set the following transformation mixture of 240 µl polyethylene glycol (50%), 32 µl LiOAc (1M), 33 µl linearized plasmid (∼ 30µg) and 5 µl ssDNA, in which the 100 µl of cell suspension were added. The mixture tube was incubated for overnight at 30ºC. The next day, the tube was heat shocked at 44ºC for 15 minutes. The cells were harvested and washed with YPD and then resuspended in 1ml. The suspension was incubated at 30ºC with shaking for 6 h. After the incubation time, the cells were harvested and spread on YPD_NAT plates. The plates were incubated at 30°C and colonies were visible after 2 days.

### Localization of Cda2, Mum3, Rrt8 and Srt1

To localize the proteins of interest, plasmids were constructed by creating fusion genes that express C-termimnal fusions to yEmRFP. First, the CIp10 vector was digested with *KpnI* and *XhoI*. Next, the gene of interest was amplified without the stop codon, using genomic DNA obtained from strain Cd1465. The yEmRFP fragment was amplified by PCR using yEpGAP-Cherry vector (56) as template. As describe above, the three fragments were fused by Gibson assembly. The plasmids were linearized by digestion with *Nco*I and transformed into the nourseothricin-sensitive mutants by lithium acetate transformation method.

### Calcofluor white (CFW)/Eosin Y staining

Chlamydospores were collected and washed with 1 ml McIlvaine’s buffer (0.2 M Na_2_HPO_4_/0.1 M citric acid [pH 6.0]) followed by staining with 30 µl Eosin Y disodium salt (Sigma) (5 mg/ml) in 500 µl McIlvaine’s buffer for 10 min at room temperature in the dark. Chlamydospores were then washed twice in McIlvaine’s buffer to remove residual dye and resuspended in 200 µl McIlvaine’s buffer. One microliter of a 1 mg/ml Calcofluor White solution (Sigma) was then added to the Eosin Y-stained cells before transfer to microscope slides. Fluorescence of Calcofluor White and Eosin Y stains was examined using DAPI and FITC filter sets, respectively.

### MDH staining of lipid droplets

To stain LDs in chlamydospore with monodansylpentane (MDH) (Abgent), chlamydospores collected as described above were washed once with 1X PBS followed by incubation in 1ml of PBS containing 100mM of MDH for 15 min in 37ºC. Chlamydospores were then washed twice with 1X PBS and examined by fluorescence microscopy using a BFP optimized filter set to visualize MDH fluorescence.

### Microscopy

All images were collected on a Zeiss Axio-Imager microscope using a Hamamatsu ER-G camera and Zen 3.0 software.

### Transmission electron microscopy

Chlamydospores were collected as described above and stained for electron microscopy using the osmium and thiocarbohydrazide staining as described previously(31). Briefly, chlamydospores were fixed by resuspension in 3% glutaraldehyde in cacodylate buffer, for 1 hr, washed once in 0.1M cacodylate buffer (pH 7.4), and then resuspended in 1% osmium tetroxide and 1% potassium ferricyanide in cacodylate buffer for 30 min at room temperature. Chlamydospores were then washed four times in dH_2_O, resuspended in 1% thiocarbohydrazide in water, and incubated for 5min at room temperature followed by one wash in dH_2_O and an additional 5 min incubation in 1% osmium tetroxide and 1% potassium ferricyanide.. The chlamydospores were then incubated in saturated uranyl acetate for 2h and dehydrated through a graded series of acetone washes. The dehydrated samples were then treated with 100% propylene oxide for 10 minutes, embedded in Epon 812, sectioned, and images were collected on an FEI BioTwin microscope at 80 kV.

## Statistics

Data presented are the mean ± SE of indicated numbers of independent samples. Statistical significance was determined with Student’s *t-*test (two-tail, heteroscedastic) using Microsoft Excel software. Differences between the analyzed samples were considered significant at *p* < 0.05.

## Acknowledgements

The authors wish to thank Neta Dean for reagents and advice, members of the Konopka and Neiman labs for helpful discussions, and Nancy Hollingsworth for comments on the manuscript. This work was supported by NIH Grant GM072854 to A.M.N. and NIH Grants R01GM116048 and R01AI047837 to J.B.K.

